# Can data mining from various internet platforms systematically accelerate detection of alien species invasions across the EU?

**DOI:** 10.64898/2026.02.06.704325

**Authors:** Simon Reynaert, Niels Billiet, Pavel Pipek, Ana Novoa, Philip Hulme, Sofie Meeus, Quentin Groom

## Abstract

Invasive alien species (IAS) expansions are increasingly impacting the biodiversity and economy of Europe. To more effectively allocate the limited resources available for their management, it is pertinent to accelerate detection of IAS spread and distribution. One largely untapped secondary data source showing much potential lies in the automated tracking of internet activity such as IAS search intensity or mentions across different internet platforms. In this study, we tested if internet activity increases systematically when IAS expand into new EU countries utilizing the combined data of 88 invasive species from various internet platforms. In total, 14 internet platforms were screened and evaluated based on their database accessibility, mined data quality and utility for systematic IAS expansion tracking. We found that the procedure to obtain researcher access to minimal data required for IAS tracking (i.e., information about location, time and place) varies widely across platforms, and is particularly difficult without incurring significant costs for many of the larger ones (X, Google and Tiktok). From the explored species, more charismatic species (i.e., mammals) overall gained more online traction than more cryptic ones (i.e., plants), though online activity of the first proved a worse representation of real-world occurrence patterns. Moreover, while the final five selected internet platforms showed increased activity surrounding the year of invasion in many of the explored invasion scenarios (particularly Wikipedia and Facebook), inconsistencies between species groups, trends per platform and the large variability in data quality currently still hampers systematic integration of such data into existing databases. We conclude that combining IAS activity data from various internet platforms shows potential to accelerate IAS expansion detection across the EU (especially for fish, crustaceans, reptiles, birds and plants). However, incorporation in automated early warning systems is currently hampered by variable data quality, limited researcher access to online data and the few open, accurate and generalizable species classification algorithms with API access.

## 1. Introduction

The cost of invasive alien species (IAS) to the global economy – ranging from damage to infrastructure and crops to depletion of natural resources (e.g., pollinator populations) and deterioration of human health (e.g., disease transmission) – is mounting every year, with Europe taking the heaviest losses (Diagne et al., 2021; Seebens et al., 2017; Soto et al., 2025). In addition, biological invasions are recognized as one of the five principal direct drivers of biodiversity loss (Roy et al., 2024) with impacts encompassing profound chemical, physical, and structural alterations of ecosystems, including changes to nutrient cycling, or fire regimes (Kumschick et al., 2024). To more effectively allocate the limited resources available for IAS management, impact mitigation and knowledge building, accelerating detection of new occurrences is becoming increasingly important (Jarić et al., 2021; Novoa et al., 2025).

Previous studies indicate that the emerging field of iEcology – exploring ecological information hidden in mentions and activity patterns of various internet platforms (e.g., social media, search engines or encyclopedias) –, shows much potential for accelerating detection of IAS outbreaks and better understanding their spread and population dynamics (Jarić et al., 2021; Novoa et al., 2025; Tomojiri & Takaya, 2025). For instance, the presence of invasive alien fish and flies was previously detected solely based on images from angler websites and Facebook groups across different EU countries (Kalous et al., 2018; Schifani & Paolinelli, 2018). Many internet platforms have also proven their utility for more systematic monitoring, such as Facebook for rare whales (Morais et al., 2021) or Flickr, Google images and Instagram for invasive plants (Allain, 2019; Cardoso et al., 2024).

Despite the abundance of potential information available online, iEcology data is fragmented (both spatially and temporarily), of highly variable quality, and usually requires thorough manual cleaning before it is suitable for ecological analysis (see e.g., issues with bots mentioned in Tomojiri & Takaya, 2025). Moreover, secondary data generated from social media mentions and internet activity will inherently be dominated by human bias (Olteanu et al., 2019). For iEcology analyses, data tends to be skewed towards larger-bodied, charismatic and easily recognizable taxa from densely populated areas with sufficient internet access (Jarić et al., 2021; Novoa et al., 2025). Thus, it is virtually impossible to use iEcology data by itself for answering many typical ecological questions relating to IAS dynamics such as, e.g., reliably estimating population size or the full extent of a species’ distribution.

Until a few years ago, monitoring efforts were mostly limited to a relatively small number of trained experts. Recently developed and widely used citizen science platforms, such as iNaturalist provide accurate data which are often useful for early detection (González-Moreno et al., 2025; López-Guillén et al., 2024). However, citizen-science platform related curation and data publishing policies often introduce lags between the time of observation and the occurrence being publicized. Any delays in data publishing may hamper rapid IAS detection (Kourantidou et al., 2022). An alternative source of real-time data, showing potential to become a useful extra component in early warning systems tracking IAS spread, lies in the monitoring and detection of changes in online activity regarding IAS (Cardoso et al., 2024; Cerri et al., 2022; Jarić et al., 2021). To our knowledge, it has not been evaluated yet to what extent iEcology datamining approaches can be applied for the systematic monitoring of IAS dynamics at a large scale (i.e., across multiple species, countries and internet platforms), and by doing so directly complement evidence-based informing of policy makers and other stakeholders regarding IAS occurrences and decisions.

Here, we investigated whether data on IAS mentions and activity from five internet platforms (Facebook, Flickr, iNaturalist, Wikipedia and YouTube) could aid in accelerating country-level detection of IAS expansions across Europe. Our overarching goals were to 1) develop automated and repeatable geolocated IAS data mining workflows and 2) evaluate whether IAS related online activity could serve as a systematic indicator for IAS expansions across Europe. Based on previous case studies (Allain, 2019; Cardoso et al., 2024; Cranswick et al., 2022; Daume, 2016; Fukano & Soga, 2019; Kalous et al., 2018; Kotowska et al., 2021; Schifani & Paolinelli, 2018), we hypothesized that IAS activity patterns across internet platforms would (i) reflect patterns in IAS occurrence data collected through active monitoring available from the Global Biodiversity Information Facility (GBIF) and the European Alien Species Information Network (EASIN) and therefore (ii) show systematic increases in IAS activity surrounding the year of invasion into new EU countries.

## 2. Methods and Materials

Before data collection, 14 online platforms were screened for their suitability as potential IAS data sources based on qualitative characteristics. These characteristics included i) the type of data (text, images or video), ii) the cost of accessing these data, iii) the geolocation precision associated with online activity, iv) the query limit for the respective application programming interfaces (APIs), v) the temporal resolution of the data, and vi) whether low-cost data mining for research purposes was authorized by the platform (Table S1).

Due to various reasons including (i) declined applications for obtaining researcher API access to perform IAS research (X, Google Ads, Google Search Researcher result API, TikTok), (ii) the lack of accurate geolocation (Reddit, Bluesky, Mastodon, eBay), and (iii) unreliable automated data mining (Google Trends; Alam & Hulme (2025)), we only incorporated data from a total of five online platforms in our final analyses. These platforms included Facebook, Flickr, iNaturalist (‘casual’ and ‘needs ID’ observations only), Wikipedia (Wikimedia) and YouTube.

The complete methodological framework included three steps (Fig. 1).In the first step, information on the 88 species included in the list of invasive alien species of the Union concern (pre-june 2025; Regulation (EU) 2016/1141) was automatically collected from Wikipedia and manually enriched using EASIN data. In a second step, we mined data through APIs about daily videos, photos, posts and activity on these species from the short-listed internet platforms mentioned above. In a third step, we processed and summarized data, conducted time series analysis and compared patterns and trends with reference datasets from the European Alien Species Information Network (EASIN) and GBIF. More details regarding all individual steps are outlined below. Data mining and processing scripts can be found in this Github repository: https://github.com/Simon-Reynaert/iEcology-IAS-miner. To protect the privacy of individuals in the public domain, only the raw count per species × country × platform data files are made available for re-analysis, with coordinates anonymized to country level and any identifiers which could potentially lead to the identification of individuals removed. Note also that the scripts for collecting and cleaning the Facebook data mining are not publicly released since Facebook data mining was performed within the Social Media Archive’s (SOMAR) secure virtual development environment at the University of Michigan (500 S State St, Ann Arbor, MI 48109, Michigan, USA)(Meta, 2025).

**Figure 1.**
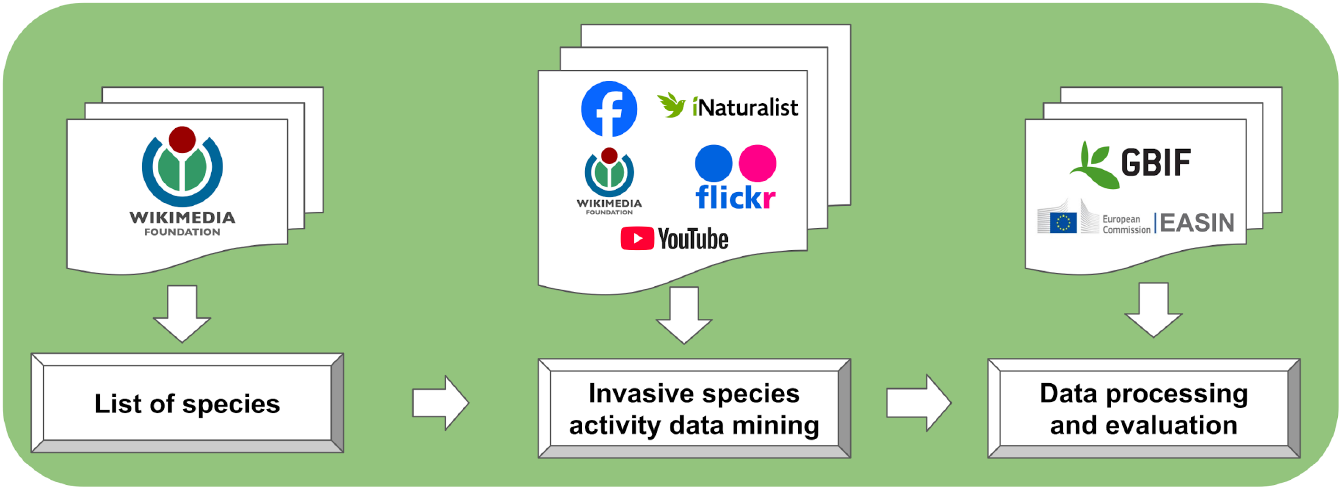
Three step methodological framework for the data collection and analysis performed in this study.

### 2.1. Data collection

All initial data collection, cleaning, anonymizing and summarizing to daily counts / activities was conducted in Python v 3.13(van Rossum, 2025). All anonymized, GDPR-compliant internet platform activity datasets, reference data from GBIF and EASIN, as well as the generated output figures for all explored EASIN invasion scenarios are available in a Zenodo repository (Link TBA).

First, the scientific names (“*Genus species*”) of 88 species included in the list of invasive alien species of Union Concern (i.e., the list pre-dating June 2025) were mined from Wikipedia and their associated Wikidata Q identifiers were stored in a data frame (see species_wikipedia_sitelinks.csv). Note that sometimes former scientific names were mined due to outdated names circulating on Wikipedia still (e.g., *Eichhornia crassipes* instead of *Pontederia crassipes*). We then collected all posts/observations/videos containing exact matches with these scientific names for countries on the European continent (Table S2) through the respective APIs of Facebook, Flickr (normal tag search), iNaturalist (only non-research grade observations) and YouTube (approximate country bounding boxes) between 01/01/2016 and 15/07/2025. Where possible, we used exact string matching within the API call to only fetch records containing the full scientific name, thus limiting collection of false positives. However, in cases where the API lacked such functionality (e.g., Facebook and YouTube), we performed post-hoc filtering to include only those posts that mentioned the full species name. For Facebook, posts were collected querying all posts containing the scientific names using the surface_countries parameter in the API to retrieve posts per country. For Flickr, we collected photos that had the species name in one of the tags such as title, description, etc. (normal tags in API call). Given every posted image on Flickr gets a new ID, we had to filter out ‘repeat’ observations made at approximately the same time and location to only count them once, in case someone had posted multiple pictures of the same observation event (i.e., one post containing multiple pictures). To collect geolocated videos from YouTube we specified a circle radius surrounding a single point coordinate for every European country in our queries, often splitting countries into multiple circles to ensure proper coverage (Table S3).

Next, we collected IAS activity data from Wikipedia. Given it is not social media (i.e., not built around posting videos/images/text), this process was slightly different. For Wikipedia, we collected daily pageview estimates over time as an indicator of platform activity, where possible by specifying “user” in the API endpoint URL to minimize false matches related to bot searches. Wikipedia daily pageviews were collected in two ways, including both geolocated pageviews (for pages with > 90 pageviews / day) and language-based pageviews (all daily pageviews except if only the editor accessed the page). Geo-located pageview data was collected by downloading historical pageview data (from 09/02/2017 until 15/07/2025) from the wikidata pageview repository (Wikimedia, 2025a) utilizing the previously mined Q identifiers to filter out pageviews related to individual species. Given the greater detection capacity (also pageviews < 90 / day) and the generally good correlations with geolocated pageview data (Fig. S1), we decided to also include pageviews that were collected through EU language filtering. To this end, we collected data from all pages linked to the wiki Q-identifier that existed in the Wikidata database and performed language to country mapping based on plausible matches (see 2.2). The accuracy of this method is known to be language dependent (e.g., we assumed pageviews to the Finnish page of invasive species primarily reflected pageviews originating from Finland, whereas this correlation is very poor for pageviews to the English page; Fig. S1)(Vardi et al., 2021). While some noise is introduced on top of geolocated pageview volumes for privacy protection reasons (Wikimedia, 2025b), we believe the effect of this on overall activity trends over time to be minimal, which was further confirmed by strong correlations with the vast majority of language based pageviews (Fig. S1).

Finally, for collection of reference and validation datasets, we directly received a dataset from the European Alien Species Information Network (EASIN) containing the year in which each species was first recorded per country for the 85 species on the Union list of Concern that were already present in Europe between 2016 and 2025. Furthermore, we collected all observations across all EU countries for the selected species from GBIF through the API and summarized them to daily counts per species and country between 01/01/2016 and 15/07/2025. After collection and initial cleaning of activity data, the daily number of activities/observations/posts per species × country combination were counted and saved into .csv files ready for analysis for all platforms except Facebook. Due to relatively small post volumes and concerns regarding user’s privacy, Facebook posts were counted and grouped per month and normalized by dividing the monthly number of posts by the maximum number of posts retrieved across the entire time series.

### 2.2. Calculations and statistical analysis

After initial data collection and cleaning which resulted in activities / observations / post counts for all queried species × country combinations across the various internet platforms, all further analyses were performed in R v 4.4.2(R Core Team, 2024). All plots were constructed using *ggplot2* (Wickham, 2016). Models were constructed with the packages *nlme* (Pinheiro et al., 2025), *lme4* (Bates et al., 2025) and *glmmTMB* (Brooks et al., 2025) and 2-by-2 differences further explored using post-hoc Tukey tests from the *emmeans* (Lenth et al., 2025) package to correct for multiple testing.

First, Wikipedia language-based pageviews were mapped with plausible candidate countries from which we assumed the majority of pageviews had originated (see also 2.1) (Vardi et al., 2021). All activities were summed per month, binning observations to increase signal detection strength while maintaining enough temporal resolution to allow detection of phenological patterns in the data. Datasets were then joined together, species synonyms matched and monthly counts used to calculate ‘popularity’, defined as the number of unique months with non-zero activity over the entire period of data collection (01/01/2016 - 15/7/2025). Differences in popularity were then compared by performing ANOVA on a GLMM (family = “negative binomial”, link = “log”) including country, taxon, habitat and internet platform as fixed effects and species nested within taxon as random intercept.

Next, we used this binned monthly data to compare individual internet platform trends with monthly GBIF occurrence data by performing Spearman rank correlations for all queried species × country combinations on the raw data and standardizing data from all platforms between 0 and 1 by dividing every monthly data point by the maximum monthly activity / post count /observation value across the entire time period. The differences in Spearman’s rank correlations were further explored by performing ANOVA on a GLMM (family = “gaussian”, link = “identity”) including country, taxon and internet platform as fixed effects and species nested within taxon as random intercept.

Finally, time series analysis was performed to detect anomalous increases in platform activity surrounding the year of reported invasion in the EASIN data for a total of 112 species × country combinations (i.e., recorded EU IAS invasion scenarios) where new invasions had occurred for the given set of species between 2016 and 2025. This was done for all raw monthly activity data for platforms individually (i.e., all 112 invasion scenarios were explored per platform) as well as for the lumped monthly sums of normalized activities across all platforms.

To assess if activity increased over time for each species on any platform for every European country, every species × country combination where a platform had sufficient data (at least one month with non-zero activity and available data for three separate months) had its counts summed per month to perform anomaly detection with the package *anomalize* (Dancho, 2023) (method = “twitter”, frequency = “auto” and trend = “auto” in the time series decomposition step; method = “gesd” with alpha = 0.25 in the anomaly detection step). The “twitter” method performs seasonal time series decomposition by piecewise medians and was specifically designed for the analysis of social media data (Dancho, 2023). Then, for all 112 EU species × country combinations with known invasions between 2016 - 2025, we calculated if a positive anomaly was present within a two year window surrounding the 1st of January of the EASIN reported year of invasion (i.e., the earliest possible invasion date +/-1 year) and if so, how many days there was between the closest increase in anomalous platform activity and that earliest possible invasion date. In other words, we detected the presence and nature of the shortest ‘lead’ or ‘lag’ expressed in number of days between the earliest possible invasion date (1st of January of the EASIN reported year of invasion) and anomalous increase in platform activity within a two year window surrounding the earliest possible invasion date. The proportion of detected anomalous activity increases during this 2-year invasion window, as well as the shortest lead and lag lengths with the earliest possible invasion date were then compared between taxa, platforms and countries using ANOVA on a GLMM (family = “gaussian”, link = “identity”) including country, taxon, habitat and internet platform as fixed effects and species as random intercept.

For the lumped data activity increase detection, normalized activity trends were summed on a monthly basis across all platforms and their behaviour characterized during the 2-year invasion window using Generalized Additive Models (GAM)(bs = “tprs”, k = 9)(Wood, 2015) to verify if combining activity data from all internet platforms improved predictability of known IAS invasions. More specifically, for every of the 112 individual known EU invasion scenarios between 2016 and 2025, we detected regions in the summed time series where either the first or second derivative showed a positive increase greater than a relative threshold (based on the total variation in observations) as well as where variance increased significantly. Per time series, only the region exhibiting the first region with a strong positive first derivative was retained. This approach prioritizes maximal instantaneous activity increases over sustained or accelerating trends, as we consider the steepest rate of increase to be the clearest and most important indicator of an immediate, anomalous activity spike. For every invasion scenario out of the 112 possible, we then verified if this region with the strongest slope and variance increases (if any) corresponded with the two-year invasion window surrounding the first of January of the EASIN reported year of invasion. Based on this ruleset, detections were ultimately classified as true positive (strong statistical signal of increased internet activity detected during invasion window), false positive (strong statistical signal of increased internet activity detected outside of the invasion window) or false negative (no significant change in variance together with first or second derivative detected across the entire timeseries).

## 3. Results

### 3.1. Mammals are the most popular species online in the EU

Our data mining exercise returned a total of 183 247 months with some online activity for the 88 species of union concern across all internet platform and country combinations (excluding the reference GBIF data). Popularity varied between platforms and was clearly skewed towards a few platforms with Wikipedia being the most popular platform on average across all unique species × country combinations (Fig. 2a; P < 0.001, Table S4). Moreover, in line with our expectations, Mammals were more popular on average across all platforms and countries compared to plants (Fig. 2b; P = 0.01; Table S5). These trends were also reflected in the rankings of individual species, with more mammals and fewer plants coming out on average as the most popular across all internet platform × country combinations on average (Fig. S2 & S3). On average, the investigated species gained much more traction online in West-European countries (Fig. S4; P < 0.001; Table S5).

**Figure 2.**
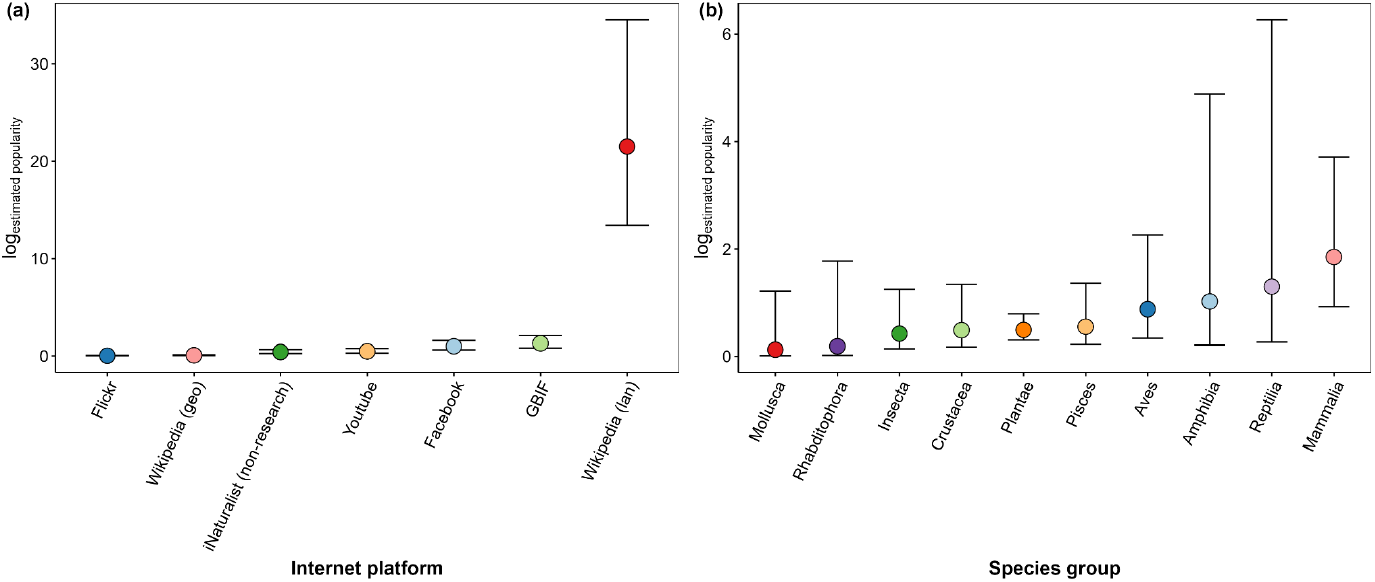
Estimated mean popularity per a) internet platform and b) species group (excl. GBIF). Error bars indicate ± 95% CI on the modelled mean per species. Wikipedia (geo) and Wikipedia (lan) refer to geolocated and language-filtered Wikipedia pageviews.

### 3.2. Internet activity of less popular species better reflects real-world GBIF occurrences across the EU

When verifying if internet activity trends reflected country-level occurrence patterns on GBIF across all queried species and country combinations, we observed that also here Wikipedia language filtered pageviews performed best (Fig. 3a; P < 0.001; Table S6), despite on average relatively small Spearman’s rho (ρ) values and large variation across all species × country observations for all platforms (Fig. 3a). Interestingly, patterns in iNaturalist non-research grade observations correlated the second best with GBIF occurrence data on average (Fig. 3a; P < 0.001; Table S6). Moreover, despite Mammalia being amongst some of the most popular species groups across internet platforms (Fig. 2b), their activity patterns aligned rather poorly with GBIF observations on average (Fig. 3b). In fact, insect activity patterns resembled most closely the available GBIF occurrence data on average – though driven by one species (*Vespa velutina*) –, and the per species correlation data also showed that 8 out of the 10 species with strongest correlations on average were Plantae, indicating that their internet activity patterns most closely resembled real-world country-level occurrences across all available data (Fig. S5). These per species rankings also indicated a tendency for internet activity to reflect trends in GBIF occurrences poorly for mammals and comparatively strongly for plants (Fig. S5), contrasting with the popularity trends (Fig. 2b). However, the per country trends were generally comparable to their popularity (Fig. S6; P < 0.001; Table S6).

**Figure 3.**
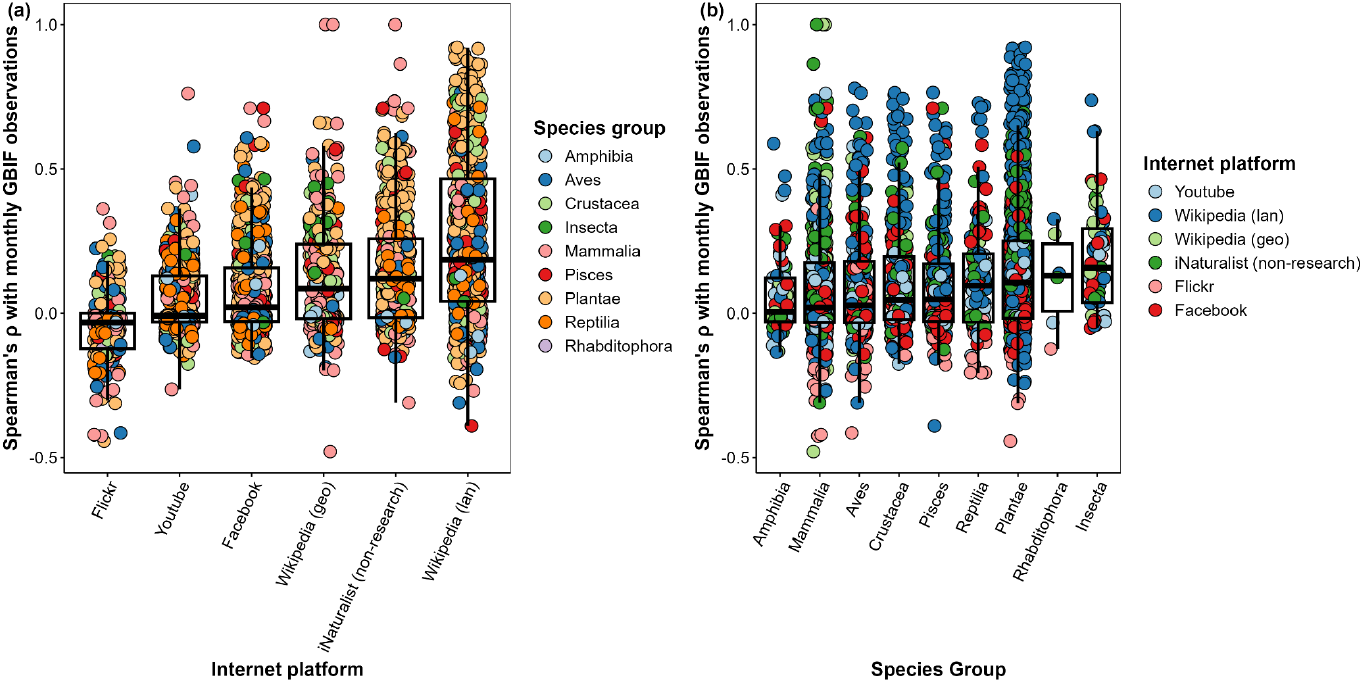
Spearman’s rho (ρ) values and boxplots of correlations between monthly summed GBIF observations and unique species × country combinations a) per platform and b) per species group. Observation points are jittered for improved clarity. Wikipedia (geo) and Wikipedia (lan) refer to geolocated and language-filtered Wikipedia pageviews.

### 3.3. Varying capacity for systematic detection of invasions from EU internet activity across species groups

Next, we explored to what extent internet activity increased during the 2-year invasion window surrounding the 112 reported EASIN invasions of 85 IAS from the list of invasive alien species of Union concern since 2016 (see e.g., Fig. S7 for the EU invasion timeline of *Pontederia crassipes*). Out of all potential species × platform combinations (excl. GBIF) where the queries returned sufficient data to be analyzed statistically, 45% (N = 275) showed an anomalous increase in online activity in the 2-year window surrounding the EASIN reported year of invasion, with proportionally most anomalies detected within (geolocated) Wikipedia and Facebook activity (60%, N = 25 and 56%, N = 81; Fig. 4a) and the least for iNaturalist non-research grade observations (26%, N = 43; Fig. 4a). Despite large variation between observations across species and countries, the closest recorded anomalous increases in internet platform activity fell on average within the year of the first record for all platforms except iNaturalist (DOY = 38 ± 178 (SD); Fig. 4b), with the reference GBIF dataset unsurprisingly being the most sensitive regarding detection of anomalous activity increases surrounding the year of invasion (69%, N = 78; Fig. 4a,b). However, anomalous data points were often also detected outside of the invasion window, leading to high false positive rates in many invasion scenarios (Fig. S8).

**Figure 4.**
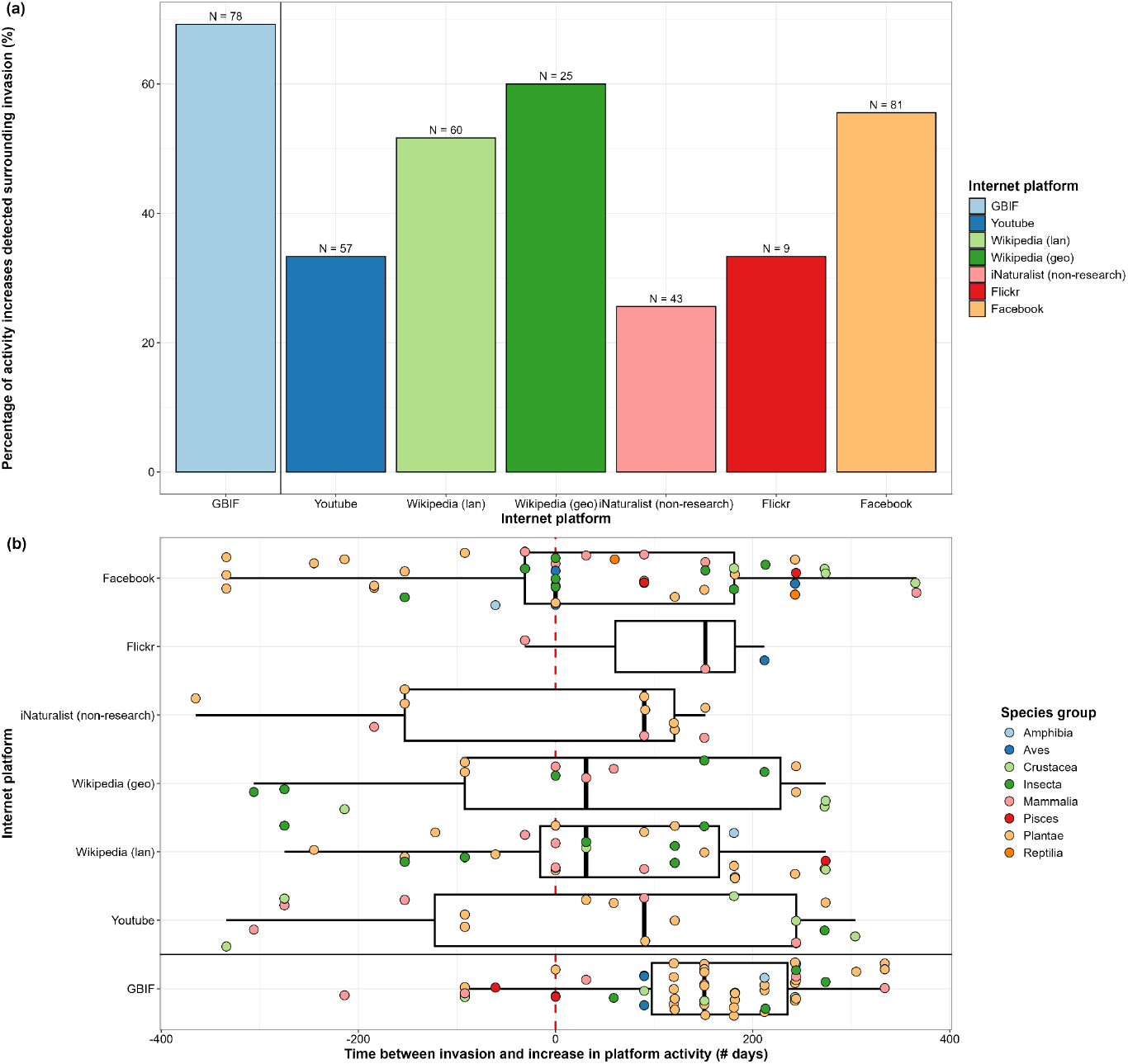
Anomaly detection data a) expressed as percentage of detections per platform and b) expressed as leads or lags from the earliest possible invasion moment in number of days per platform. In a) the total number of potential species × country combinations with sufficient data for anomaly detection per platform are indicated as N above each bar. The vertical red dashed line in b) indicates the first of January of the EASIN reported year of invasion. Observations are jittered for improved clarity. Wikipedia (geo) and Wikipedia (lan) refer to geolocated and language-filtered Wikipedia pageviews.

To tackle the issue of anomalous internet activity spikes outside of invasion windows inflating false positive rates on a per platform time series basis (see e.g., Fig. S9), we pooled internet platform data together and performed activity increase detection on the cumulative sum of normalized monthly platform activity data for each species × country combination. When pooling data and looking at combined slope and variance increases rather than isolated anomalies, overall true positive detection rate was 45% (N = 112; Fig. 5a, b). However, while the first strong signal of increasing internet activity overlapped with the invasion window for many species × country combinations, our detection algorithm still returned 31% false positives (N = 112; Fig. 5b), due to comparatively stronger activity increase signals in non-target regions of the respective time series. In addition, for 24% of the explored scenarios, no strong activity increases were detected throughout the entire time series(N = 112; Fig. 5b). Among taxonomic groups, true positive detection rates (TPR) were greater than false positive rates for Aves (TPR: 80%, N = 5; Fig. S7), Pisces (TPR: 75%, N = 4; Fig. S7), Reptilia (TPR: 67%, N = 3; Fig. S7), Crustacea (TPR: 57%, N = 14; Fig. S7) and Plantae (TPR: 25%, N = 64; Fig.S7), equal to false positive rates for mammals (TPR: 36%, N = 11; Fig. S7) but smaller compared to false positive rates for Insecta (TPR: 10%, N = 10; Fig. S7).

**Figure 5.**
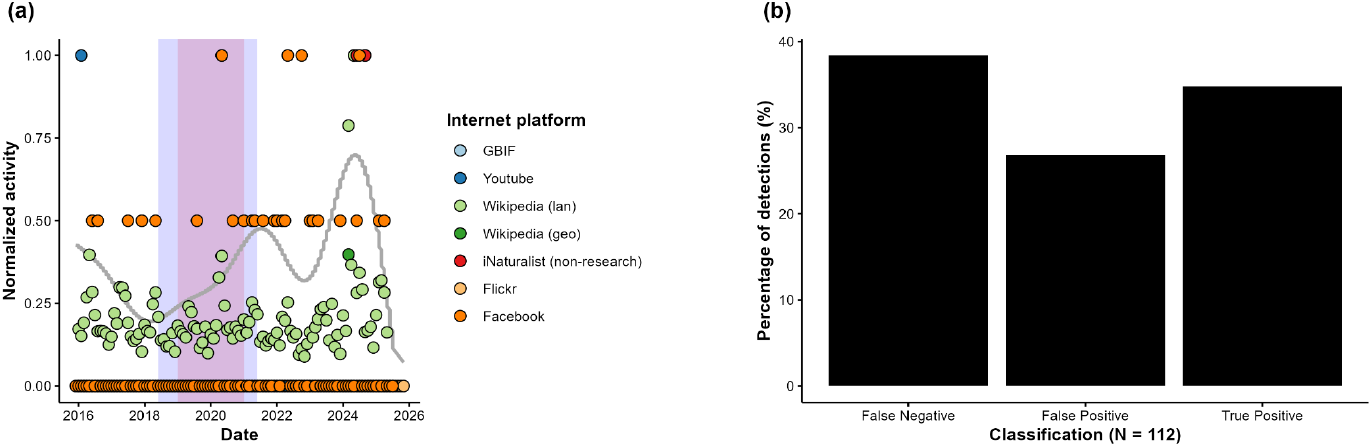
Example of a) the mean fitted GAM predictions (grey line) on the sum of normalized monthly activities for the combined internet platform activity data (excl. GBIF) of *Pontederia crassipes* in Poland (see also Fig. S4). The red box outlines the 2-year invasion window (reported EASIN invasion year in 2020) and the purple box indicates the first region of strong activity increase in the summed normalized data timeseries based on our pre-defined threshold. GBIF data was not used in the models but is shown for illustrative purposes. In b) the sensitivity of the detection algorithm is summarized across the total of 112 unique EASIN reported invasion cases of species of the Union list of concern between 2016 and 2025.

## 4. Discussion

Tracking online activity (mentions / searches) regarding IAS across internet platforms shows much potential to accelerate detection of novel species invasions (Daume, 2016; Jarić et al., 2021). We explored whether combining geolocated activity data (posts, mentions, videos, pageviews, searches) on IAS from different internet platforms (Wikipedia, Youtube, iNaturalist, Facebook, Flickr) could be used as a means to systematically identify new invasions across the EU. Out of all platforms, Wikipedia and Facebook were the most popular (i.e., had the most unique months with non-zero activity) for the explored species × country combinations. However, despite an abundance of IAS activity data across various internet platforms, data availability and quality varied strongly across species and platforms, with a tendency for bias towards sustained interest for more active and often considered charismatic species groups (i.e., mammals) but more accurate reflections of real-world occurrence patterns for less popular species such as plants. When exploring recent early invasion scenarios across the EU, we found that Wikipedia followed by Facebook were the most useful platforms for detecting new species invasions, detecting anomalous activities in about 60% of cases with sufficient data. However, even when pooling data together for all platforms, false positives were still flagged in approximately one third of the invasion scenarios, with the highest likelihood of detecting true IAS invasions (> 50% of cases with clear activity signal) from internet activity data for birds, fish, reptiles and crustaceans versus the lowest for plants, mammals, insects and amphibians (< 50% of cases with clear activity signal).

Contrary to previous case studies (see e.g., Jarić et al. (2021)) and our first hypothesis, correlations between internet platform activity data and real-world GBIF occurrences per country were overall relatively weak for all platforms. Despite that some platforms such as Wikipedia, iNaturalist and Facebook showed a tendency to correlate better with real-world GBIF occurrences – likely related to persistent identifiers aggregating activity data per species topic rather than individual keywords or tags such as in other platforms –, their mined data was still of highly variable quality. Some species did show strong correlations with real-world occurrences across platforms and countries (e.g., *Microstegium vimineum* or *Vespa velutina*), but these were exceptions rather than the rule. Nonetheless, internet activity data relating to plants showed a tendency to correlate better with occurrences mined from GBIF compared to mammals, despite the latter being considerably more popular across all platforms. One potential explanation for this observation could be that internet activity data of more popular species are inflated with false positives, leading to less accurate representations of real-world occurrence patterns. Given these inconsistencies and the inability to manually verify observations with such big data volumes, these results highlight the urgent need for generalizable, open-source text and image based machine-learning tools that allow verification of rare invasive species identity with reasonable certainty from internet activity data, similar as to what has been already developed in more specific contexts (e.g., Cardoso et al., (2024)).

Interestingly, only three internet platforms (Flickr, Youtube and iNaturalist) allowed mining of IAS images, videos and occurrences with exact geolocated coordinates. Whilst the use of geolocated internet activity data without anonymization is problematic within the public space due to conflicts with privacy legislation (e.g., GDPR) and the use of internet platform data outside of the user-consented context (Di Minin et al., 2021; Sandbrook et al., 2021; Thompson et al., 2021; Xue,2024), the lack of real coordinates associated with most mineable iEcology observations hampers direct integration into secure databases that could be used in specific national or international contexts where research regarding environmental issues and public health (i.e., matters of ‘public interest’) outweigh privacy protection rules. Thus, the utility of using automated iEcology data harvesting workflows to collect primary species observations remains overall likely limited to only a handful of platforms.

In that regard, although our research questions fall under the assessments of risks from digital data outlined in the EU’s digital service act (EU, 2022), and that we ensured secure and anonymized data processing and publication practices, multiple platforms still rejected our researcher access API applications (X, Google Ads, Google Search) or only allowed us to extract already anonymized (i.e., to country level), normalized or summarized activity data points (Facebook; Vardi et al. (2021)). As such, not only has internet platform data quality declined significantly over the past few years due to exceedingly greater numbers of bot accounts, users deliberately pushing misinformation and manipulative algorithms maximizing profit and political gain over content moderation (Koo et al., 2025; Ye et al., 2025), but the access to such data has also been significantly restricted (Hase et al., 2024). Two recent examples of such restrictions are directly impacting prominent iEcology and culturomics data sources. First, is the loss of researcher access to X’s data, where iEcology researchers used to have access to through the X’s API in the past (Novoa et al., 2022; Tomojiri & Takaya, 2025). Second, is the phasing out of popular Python (*Pytrends*, 2025) and R (Massicotte & Eddelbuettel, 2022) libraries for the automated mining of Google Trends data, since this feature is being integrated into the Google cloud system as a dedicated API with (currently) unknown researcher accessibility options (Google, 2025).

Our second hypothesis was confirmed to some extent. In approximately half of the explored scenarios that yielded sufficient data, there were spikes of platform activity surrounding the first report of the invasion of an IAS into a new EU country. However, a large slice of the tested scenarios did not return any useful data across platforms. In line with the popularity and the correlations with GBIF records, anomaly detections were most informative for Wikipedia, Facebook and Youtube, with Facebook returning data for most species × country combinations out of any platform including GBIF (81 out of 112), despite much of these combinations showing no spikes of activity during the actual invasion window. In addition, our results suggest that the occurrence datasets mined from GBIF should not be used as the sole data source for early warning systems: only 78 out of 112 species × country combinations yielded data and we could not capture anomalous occurrence signals after the IAS were first detected in each country in about 30 % of the cases with sufficient data. Moreover, while many platforms detected useful anomalous activity increases during the invasion windows, most internet platform activity time series also showed anomalies activities in non-target regions, inflating the number of false positives massively and limiting the use of such an approach within early warning systems. Despite these limitations, citizen science platform data mined through GBIF has shown its use for capturing first records of invasive species, albeit inconsistently across countries and taxa (González-Moreno et al., 2025).

To tackle the issue of numerous false positives, we pooled activity data together and verified if this improved invasion detection capabilities. Overall, this method indeed improved overall true positive to false positive rates, despite still returning many false negatives. As such, the likelihood of flagged anomalous activity windows reflecting real-world invasions became somewhat higher. However, the ability of our algorithm to detect real invasion windows still differed between species groups. Interestingly, despite mammals being among the most popular species across platforms, our algorithm only detected a few real invasion windows, while the processing of fish, crustaceans and reptile invasion scenarios virtually only returned true positives. This observation is in line with how well the activity data seemed to reflect real world occurrences, further confirming that internet activity data on the most popular species often offers a worse representation of real world patterns (though with exceptions such as *Vespa velutina*).

Some shortcomings of the methodology used in this study should be highlighted. First, due to reasons relating to API rate and query limits (YouTube), disproportionate amounts of false positives requiring manual verification (Flickr) as well as reasons of practical implementation within VDE’s (Facebook), most of our queries were limited to a keyword list only including the scientific names of the target species. Although this in all likelihood resulted in some missed ‘real’ IAS observations, we believe that the detected activity changes would still follow the same patterns as when including all synonyms and common names, given a single observation was enough to be classified as anomalous activity. Additionally, querying on scientific names assumes that users investigating the subject online have at least some intermediary level of knowledge about the species, increasing the likelihood of such observations actually reflecting the species of interest. Nonetheless, one of the reasons why Wikipedia returned disproportionate volumes of data compared to other platforms may be related to the fact that they work with unique identifiers linking up common names, synonyms and scientific species names within one topic, summing all activity relating to these topics together under one persistent umbrella. Given that species taxonomy changes on a regular basis and that keyword lists are per definition always finite and incomplete, the ability to extract activity data relating to a topic is a large benefit of this platform over others. Secondly, data was processed in ways that could influence the outcomes of this study. It is possible that binning observations to a monthly sum diluted important anomalous signals that occurred on a finer temporal resolution. However, binning was necessary to allow comparison between all platforms, and we are quite confident that large anomalies were still visible, given that phenological signals were also still present in the monthly time series data. Thirdly, we considered the ‘invasion window’ to be the two-year period surrounding the earliest possible invasion date as reported by EASIN, which is somewhat an arbitrary middle point. As such, it is possible that some platform activity spikes were missed if e.g., the species only invaded towards the end of the second year and activity spiked with a lag. However, expanding these detection windows would go beyond investigating whether internet platform activity increases systematically during the early stages of real-world invasions. Finally, after having explored multiple options and benchmarked our thresholds, we chose to characterize our increases by looking at the first variance increase that coincided with positive increases in the first and second derivative within our time series (similar to the method proposed in Vanderhoeven et al., 2017). While the concept of ‘invasion fronts’ is commonly used in invasion biology (Arim et al., 2006), no studies thus far have explored whether such fronts are also systematically found in iEcology data. Given the more noisy nature of iEcology data per definition (Cardoso et al., 2024; Jarić et al., 2021), these criteria may thus not be the ultimate way of detecting alien species invasions from internet platform activity data.

## 5. Conclusion

We explored whether geolocated IAS internet platform activity patterns can be used as a systematic early warning system for detecting real-world species invasion across the EU. While our results indicate that online activity patterns of some taxonomic groups (fish, crustaceans, reptiles, birds and plants) reflect real-world early invasion patterns quite well, inconsistencies across species and countries currently hamper development of systematic detection algorithms and implementation in decision tools. Moreover, given the noisy nature of iEcology data, the recent trend across internet platforms to limit researcher access to well-documented databases through automated programming interfaces (Hase et al., 2024), likely worsens sensitivity of detection of real-world invasions due to the increasingly more limited access to geolocated online observations. To improve the utility of iEcology data for informing researchers and policy makers regarding invasive species expansions across the EU, future studies should (i) aim to develop more open, accessible and generalized invasive species machine vision models, (ii) work on systematic improvement of data processing algorithms that minimize retention of false positives across platforms, and (iii) push for improved researcher access to detailed internet platform data within the context of identifying systemic risks from online data as outlined by the digital service act.

## 6. Data availability statement

The data and code associated with this study is openly available in the Zenodo repository: TBA.

## 7. Acknowledgments

This study was supported financially by the EU horizon project OneSTOP (Grant agreement ID: 101180559).

## Supplementary material

**Table S1.**
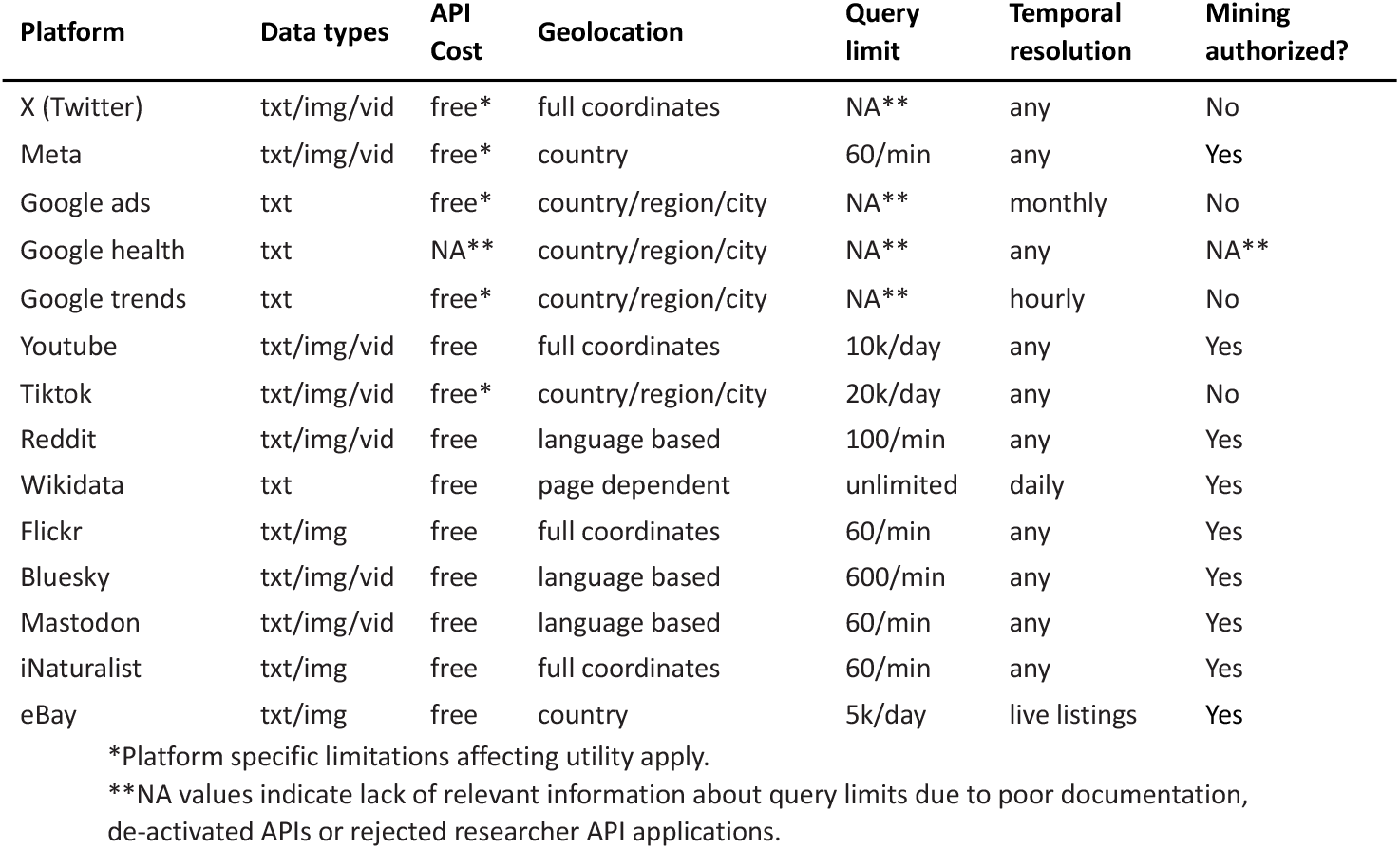
Characteristics utilized to evaluate the suitability of different internet platforms for mining iEcology data through APIs. Text or numerical values, images and videos are abbreviated by txt, img and vid, respectively.

**Table S2.** The Alpha-2 ISO codes and country names of European countries included in the API data collection queries.

| Alpa-2 ISO Code | Country name |
| --- | --- |
| AT | Austria |
| BE | Belgium |
| BG | Bulgaria |
| HR | Croatia |
| CY | Cyprus |
| CZ | Czechia |
| DK | Denmark |
| EE | Estonia |
| FI | Finland |
| FR | France |
| DE | Germany |
| GR | Greece |
| HU | Hungary |
| IE | Ireland |
| IT | Italy |
| LV | Latvia |
| LT | Lithuania |
| LU | Luxembourg |
| MT | Malta |
| NL | Netherlands |
| PL | Poland |
| PT | Portugal |
| RO | Romania |
| SK | Slovakia |
| SI | Slovenia |
| ES | Spain |
| SE | Sweden |
| IS | Iceland |
| LI | Liechtenstein |
| NO | Norway |
| CH | Switzerland |
| GB | United Kingdom |

**Table S3.**
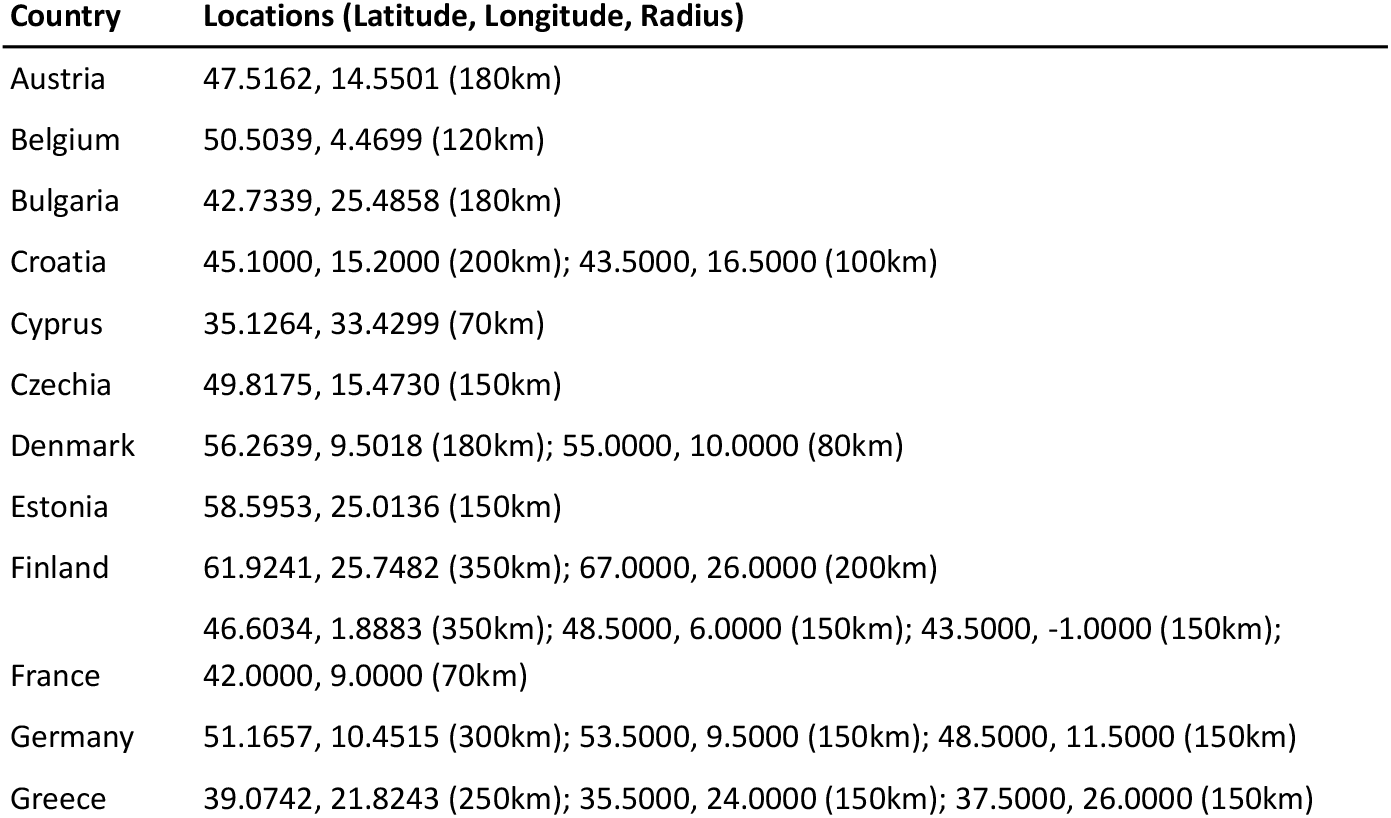
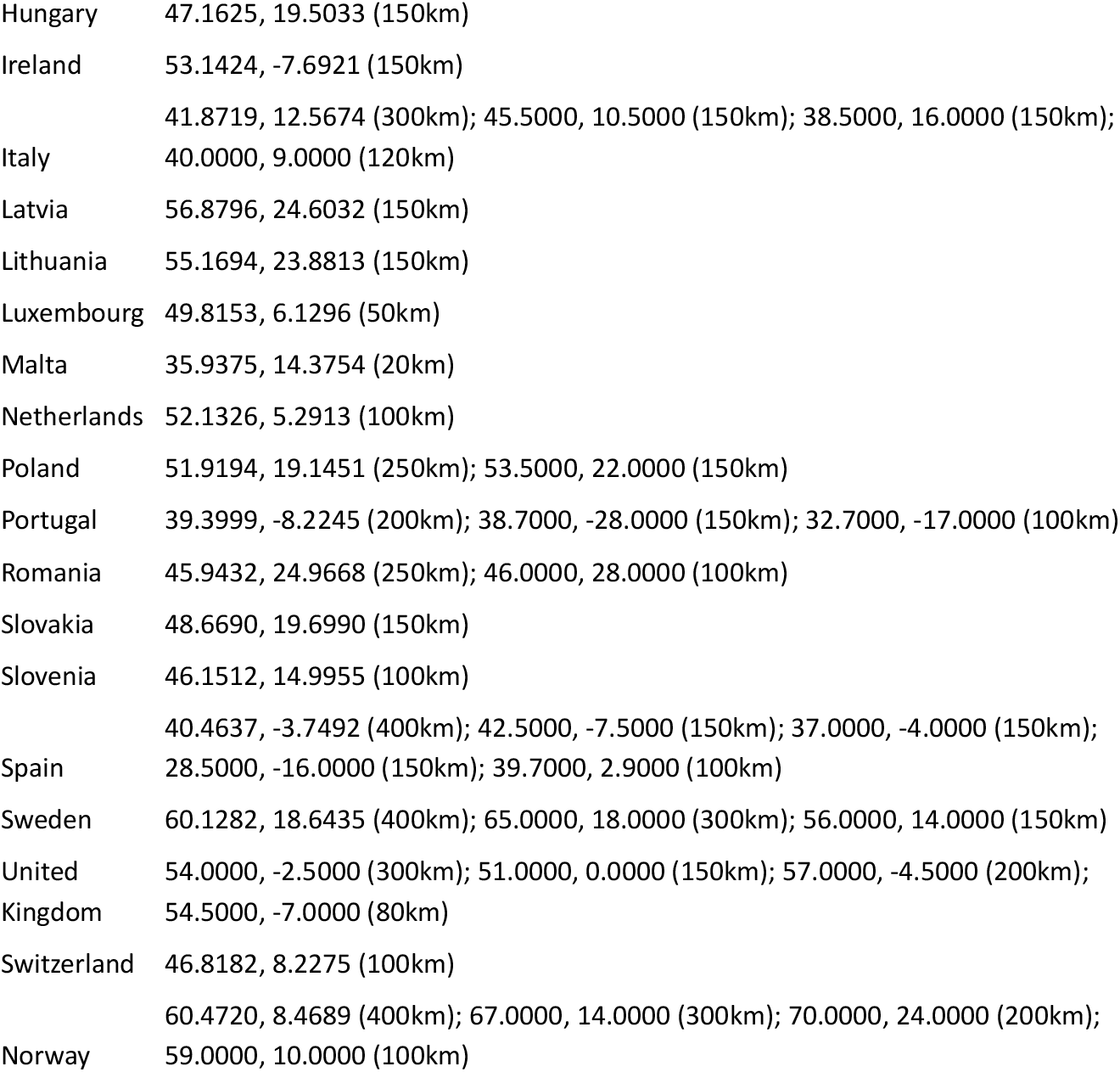
Different European countries and their approximate coverage by different circles used for the Youtube API queries.

**Figure S1.**
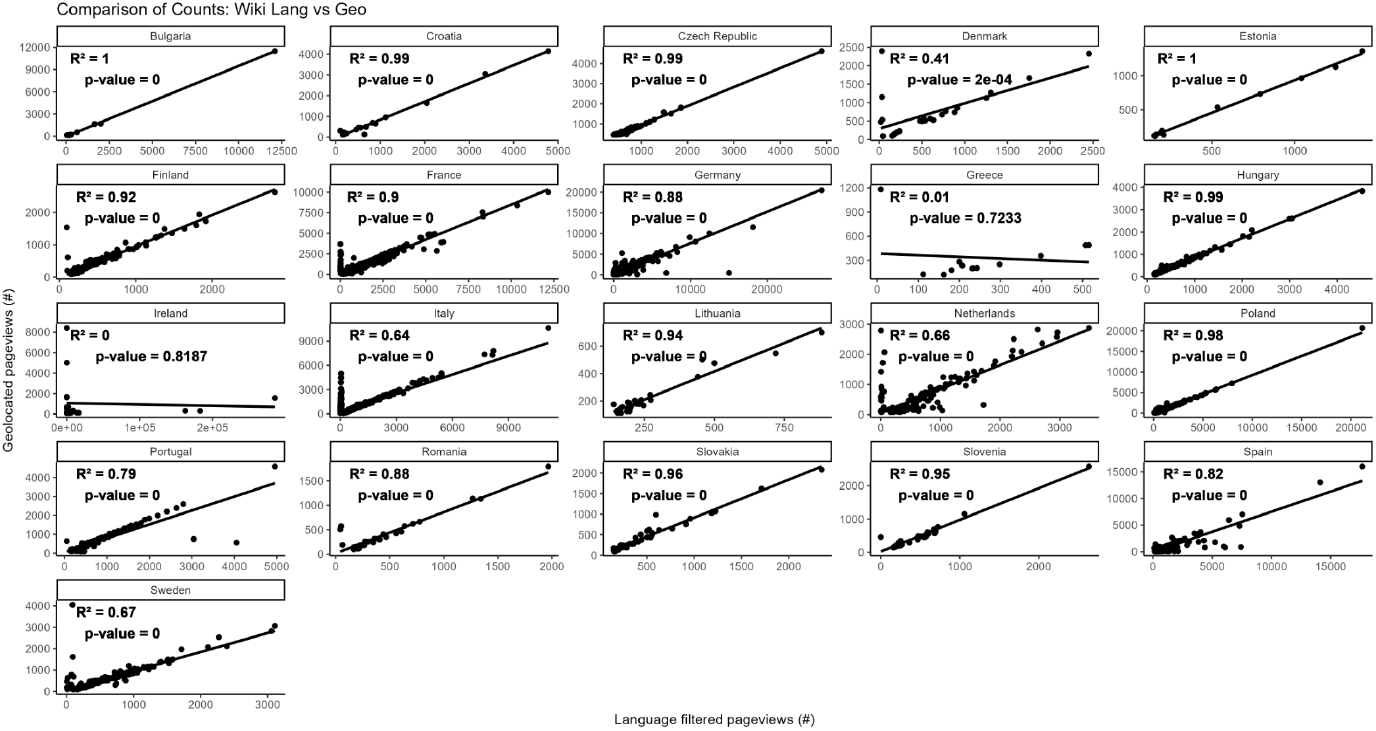
Correlations of non-NA observations between wikipedia pageviews per country based on language filtering vs direct geolocation (> 90 pageviews per day) for all species on the Union List of Concern.

**Table S4.** Type II ANOVA outcome table of the GLMM (family = negative binomial, link = ‘log’) testing for the effects of internet platform (incl. GBIF), taxon, country and habitat on popularity.

| Response variable | Fixed effect | df | Chisq | P - value |
| --- | --- | --- | --- | --- |
| Popularity | Internet platform | 6 | 7870.825 | **< 0.001** |
|  | Taxon | 9 | 16.981 | **0.049** |
|  | Country | 50 | 3647.887 | **< 0.001** |
|  | Habitat | 2 | 3.075 | 0.215 |

**Table S5.** Type II ANOVA outcome table of the GLMM (family = negative binomial, link = ‘log’) testing for the effects of internet platform (excl. GBIF), taxon, country and habitat on popularity.

| Response variable | Fixed effect | df | Chisq | P - value |
| --- | --- | --- | --- | --- |
| Popularity | Internet platform | 5 | 8229.114 | **< 0.001** |
|  | Taxon | 9 | 21.680 | **0.01** |
|  | Country | 50 | 3081.641 | **< 0.001** |
|  | Habitat | 2 | 2.641 | 0.267 |

**Figure S2.**
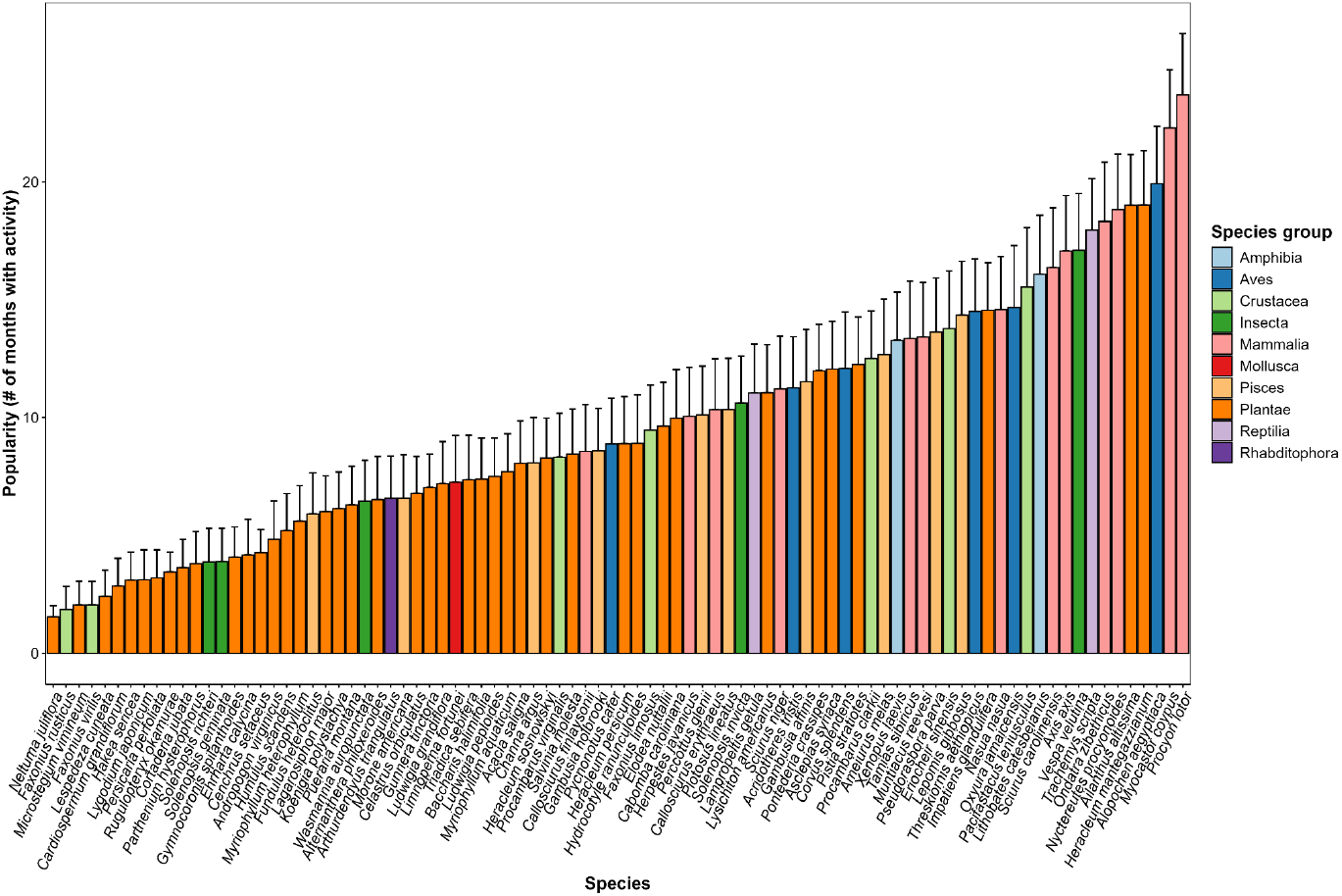
Average popularity of species across all internet platform x country combinations ranked by increasing popularity. Error bars represent +/-1 SE on the mean.

**Figure S3.**
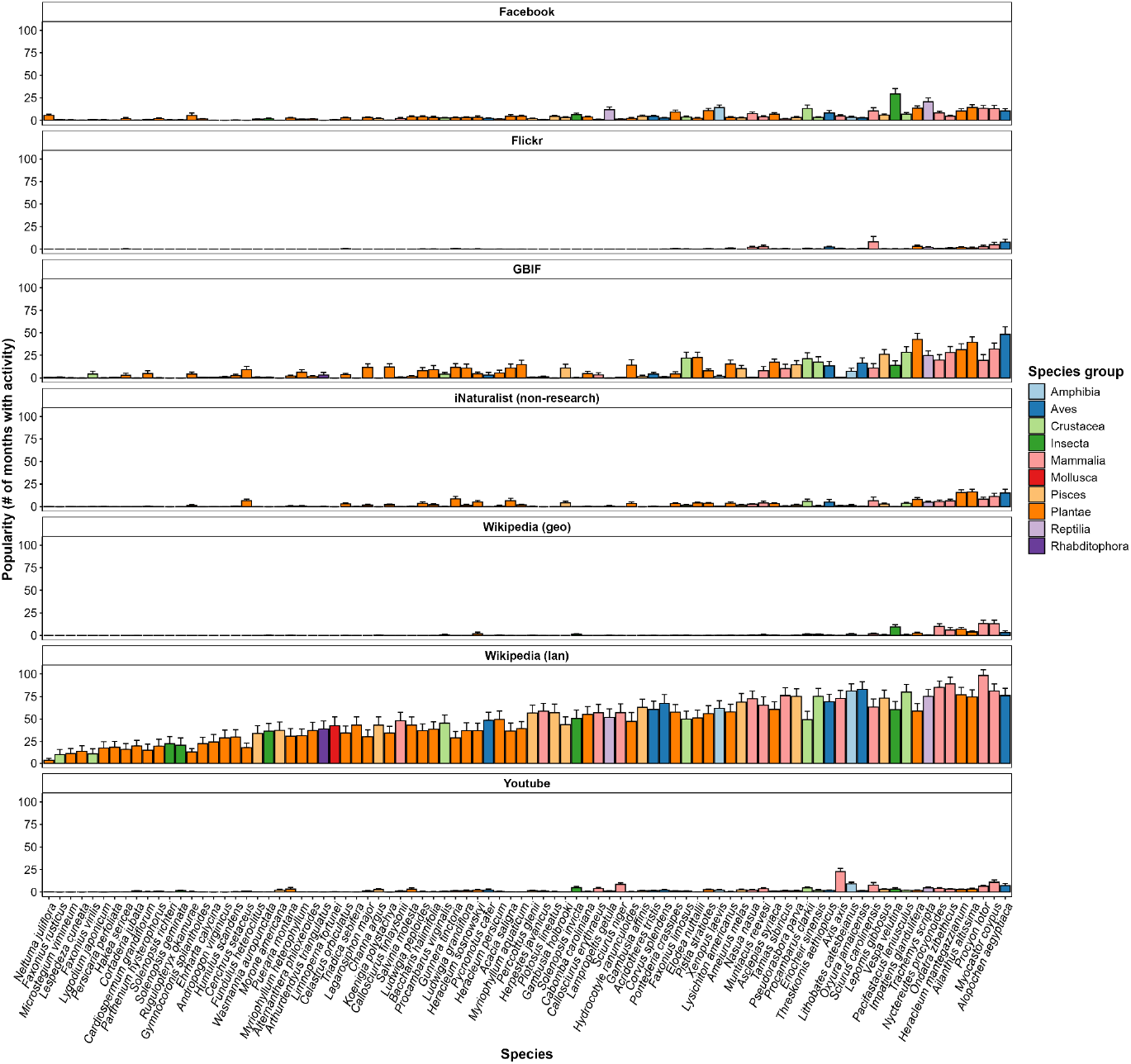
Average popularity of species per platform across all countries ranked by increasing mean popularity across all observations. Error bars represent +/-1 SE on the mean.

**Figure S4.**
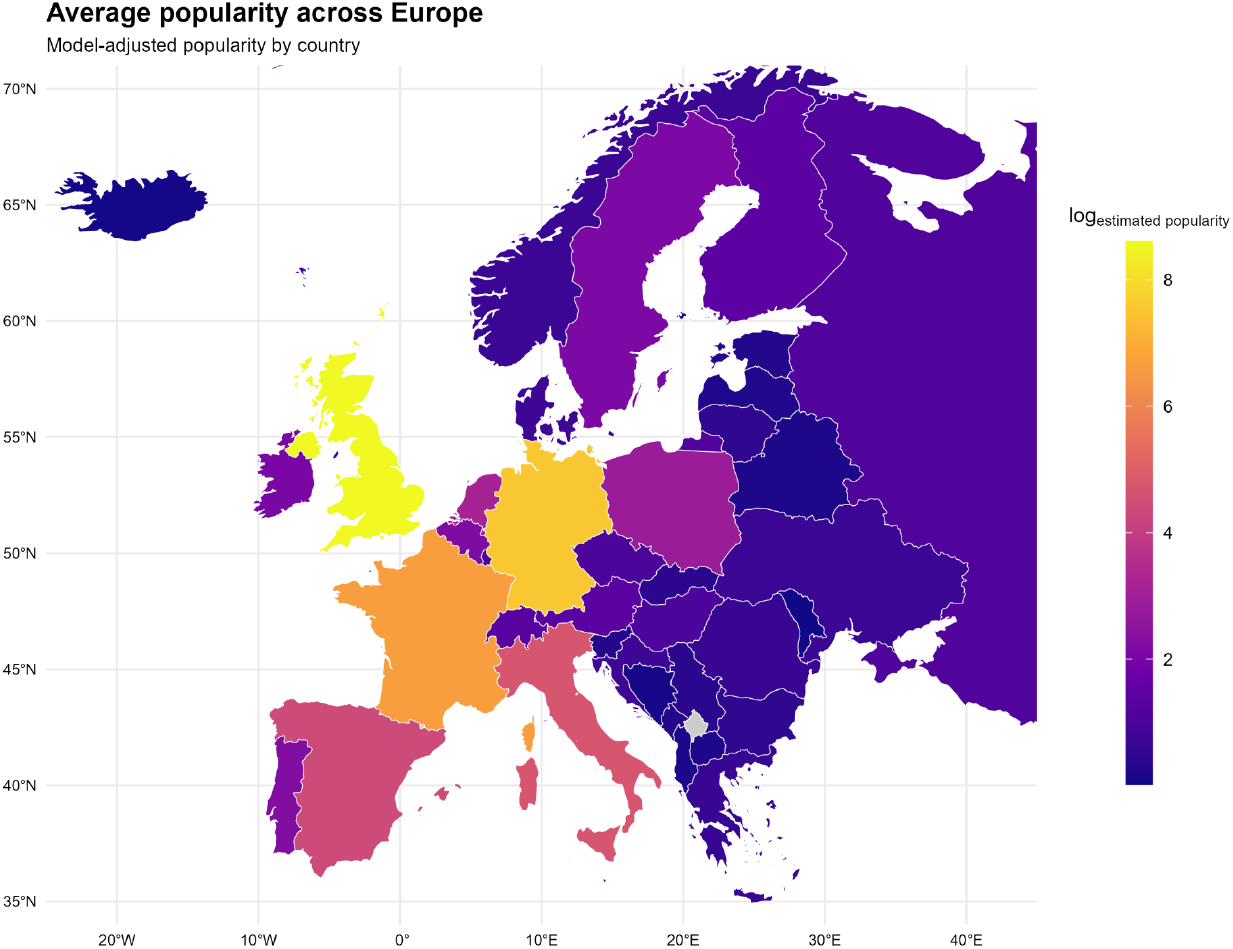
Map of modelled mean natural logarithm of popularity per country across all species.

**Table S6.**
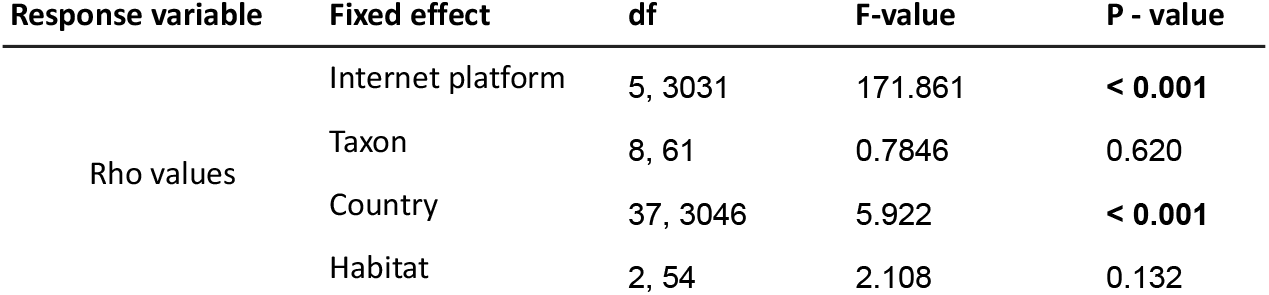
Type III ANOVA outcome table of the GLMM (family = gaussian, link = ‘identity’) testing for the effects of internet platform, taxon, country and habitat on rho value outcomes of the correlations between monthly internet platform activity data and GBIF occurrences.

| Response variable | Fixed effect | df | F-value | P - value |
| --- | --- | --- | --- | --- |
| Rho values | Internet platform | 5, 3031 | 171.861 | **< 0.001** |
|  | Taxon | 8, 61 | 0.7846 | 0.620 |
|  | Country | 37, 3046 | 5.922 | **< 0.001** |
|  | Habitat | 2, 54 | 2.108 | 0.132 |

**Figure S5.**
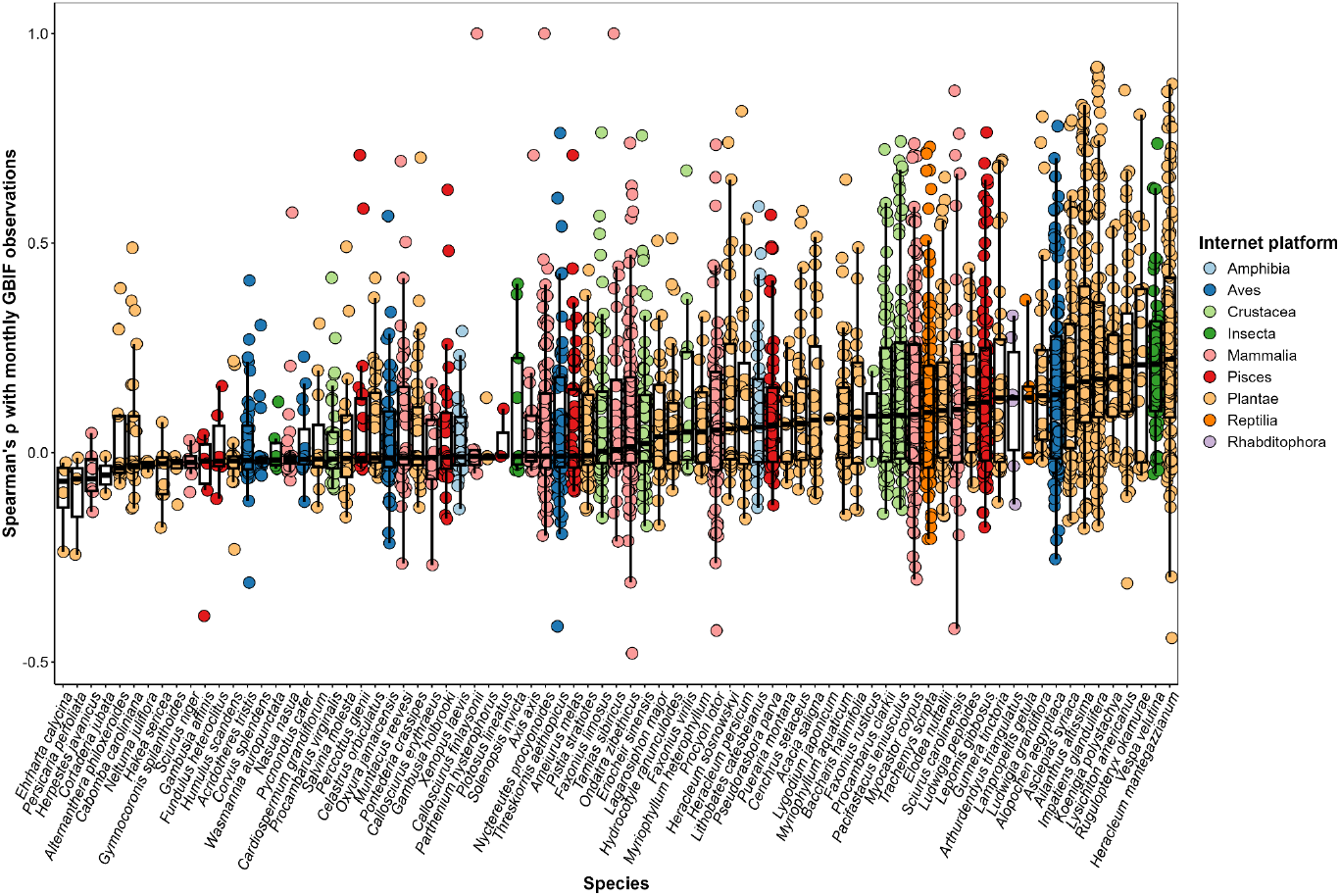
Spearman’s rho (ρ) values and boxplots of correlations between monthly summed GBIF observations and unique platform x country combinations per species.

**Figure S6.**
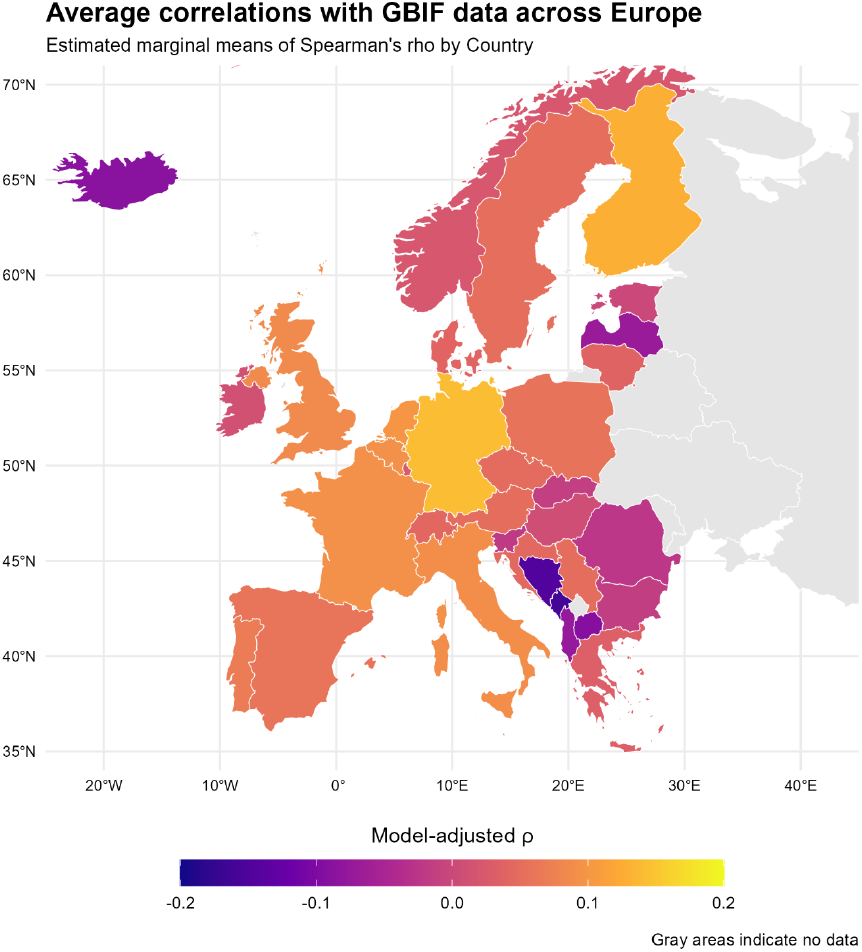
Map of Europe showing mean modelled Rho values of average monthly internet activity across all platforms and species with GBIF occurrences.

**Figure S7.**
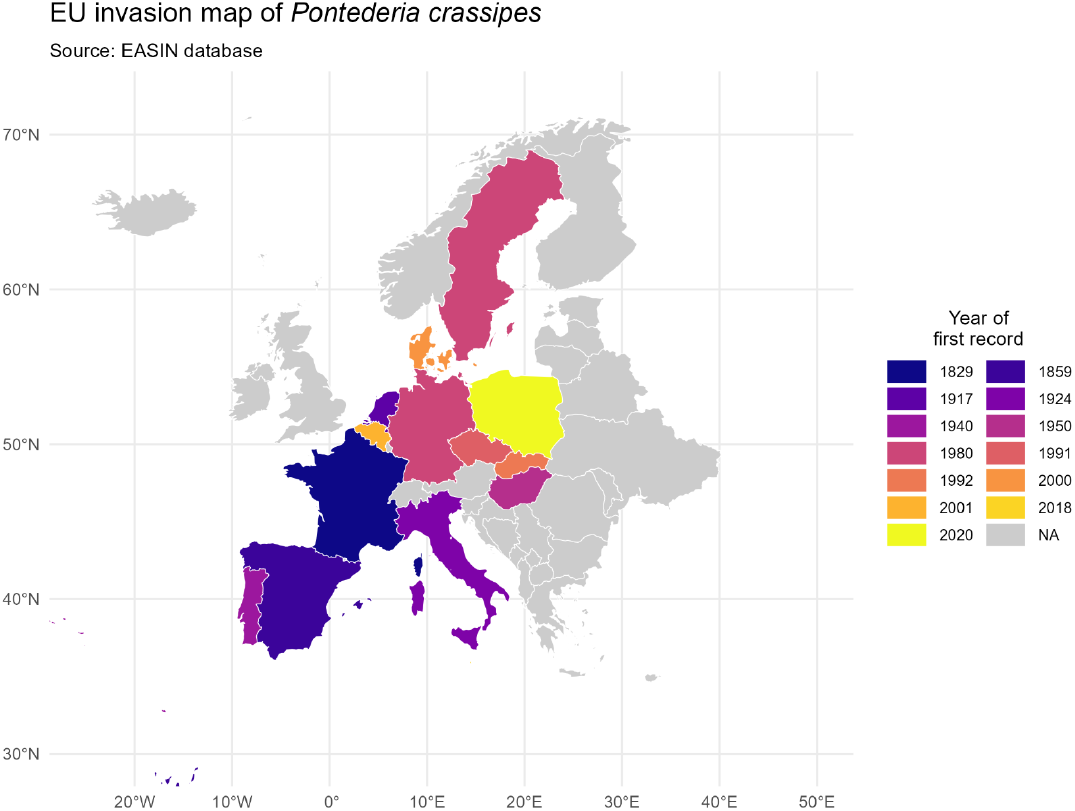
Invasion map of *Pontederia crassipes* across the EU with the EASIN reported invasion years highlighted. This species invasion in France from 2019 onwards was used as an example for the sensitivity analysis (see also section 3.3).

**Figure S8.**
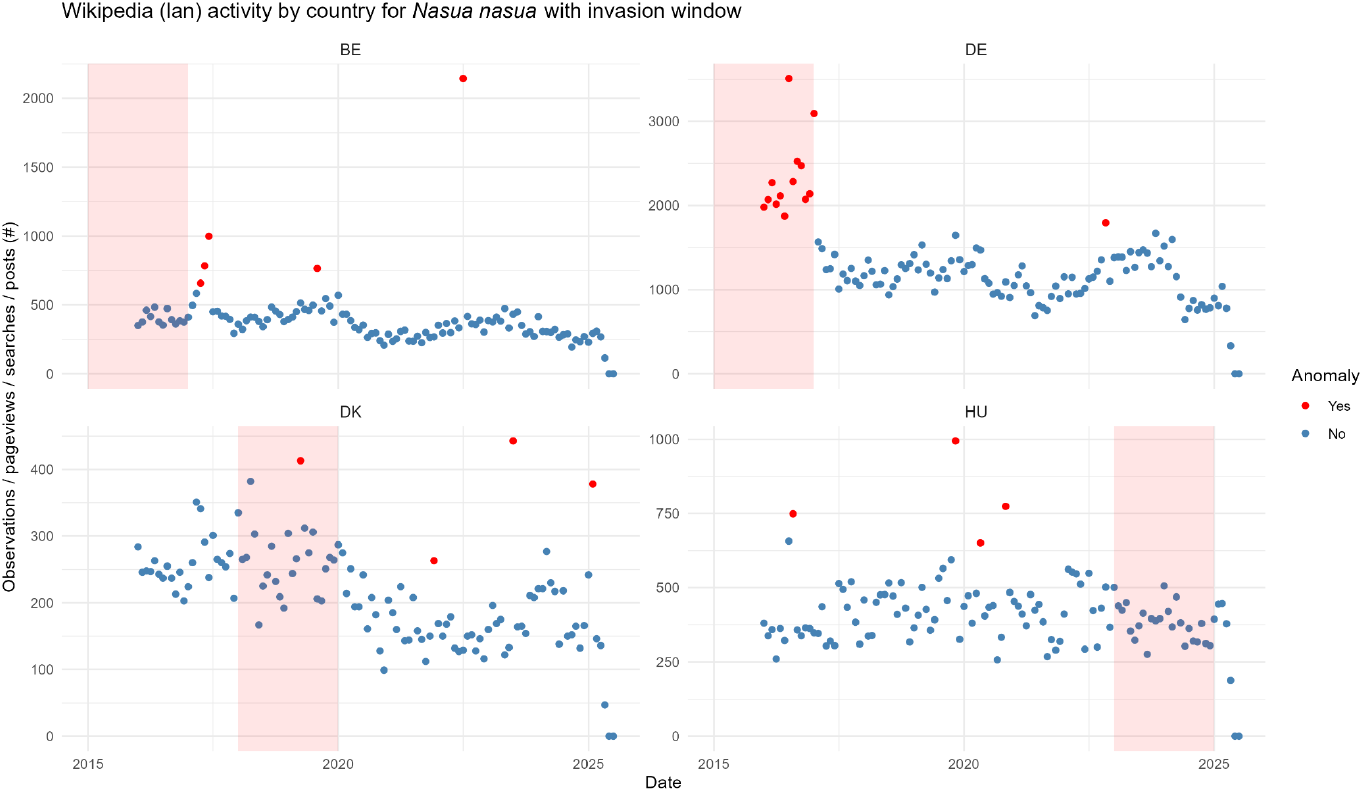
Wikipedia language filtered pageview timeseries and their anomalies highlighted in red for the 7 invasion scenario’s of *Nasua nasua* in the EU since 2016. The ISO-3166-1 alpha-2 country codes are given above the individual facets. The red box outlines the two-year invasion window including both the EASIN reported year of invasion and the year prior.

**Figure S9.**
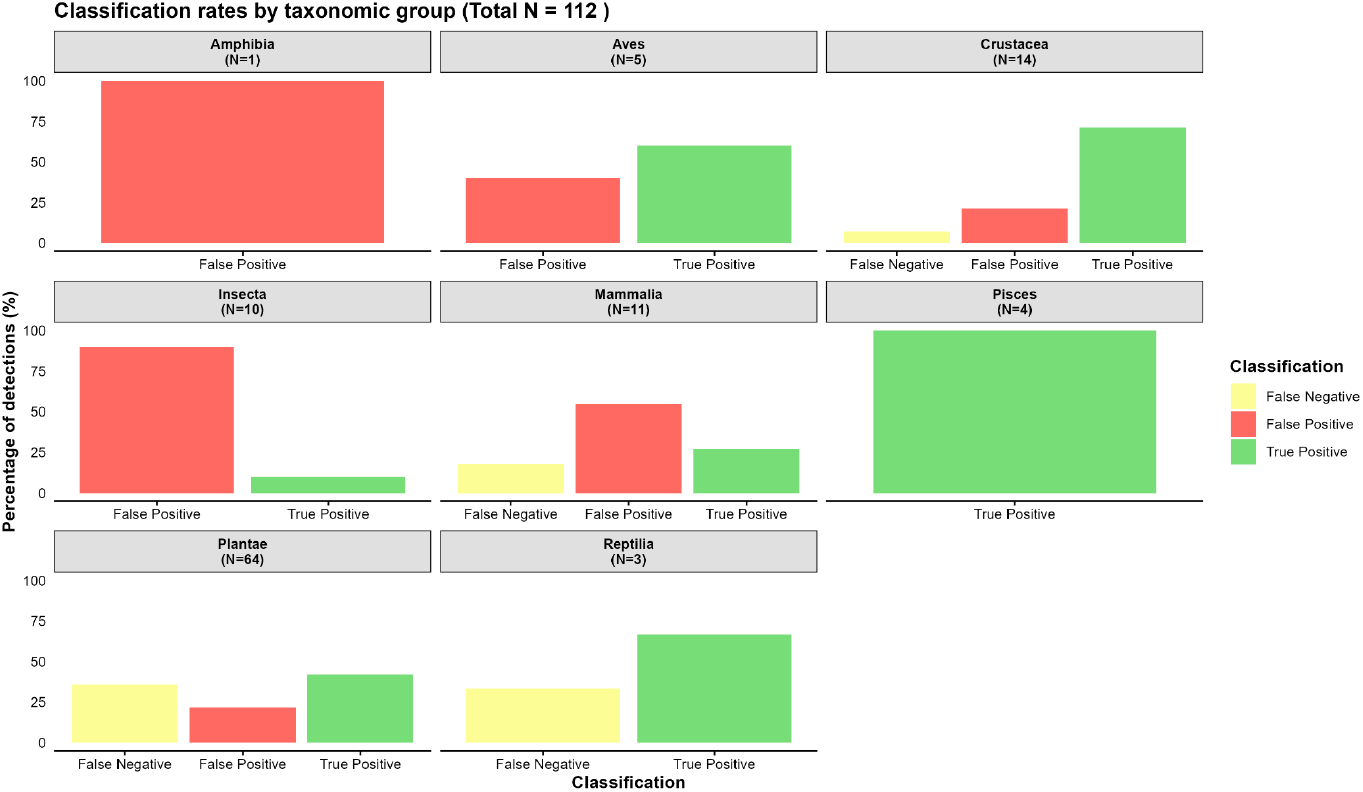
Sensitivity analysis of activity increase detection rates by taxonomic group.

